## Supplemental text, tables and figures for "Accurate pan-cancer tumor purity estimation from gene expression data"

### Supplementary

#### Training machine learning models

Linear Regression, nuSVR, Elastic Net, Lasso and Gradient Boosting models were trained using the scikit-learn Python library (Supplementary, Python code). Logit Regression was built as a modified version of Linear Regression from scikit-learn, Fully Connected Neural Net with variable layer size was built using Keras module of the Tensorflow Python library (Supplementary, Python code). All the hyperparameters not explicitly defined in the model call functions (e.g. alphas for Lasso) or in the hyperparameter search functions (e.g. HalvingGridSearchCV), were allowed to either be chosen by the in-built hyperparameter selection procedure or be used at their default values.

#### Overview of existing transcriptomics-based approaches capable of predicting tumor purity

Purity prediction from gene expression matrix-like data is an example of a well-studied task of *computational deconvolution* (1). Over the last few years, an abundance of techniques using bulk gene expression data as an input has been developed. However, only some of the existing methods are capable of performing deconvolution based solely on the input data matrix and a set of constraints included in the algorithm or in the reference data. Below is a brief overview of some widely used transcriptomics-based methods that are capable of predicting cancer cell proportions, given only gene expression matrix and, sometimes, other data included with the software.

CIBSERSORTx (2) works by first building a signature matrix of the most significant genes in pre-specified cell types and then, assuming a linear mixing model, performing $\nu$-support vector regression to infer the proportions of cell types in each sample.[13] Its most prominent limitation is that CIBSERSORTx relies on the signature matrix created from single-cell data and thus introduces certain bias into the deconvolution of samples from other experiments. Although the authors provide the signature matrix for some cancer types, for most of the new data the matrix has to be reconstructed de novo (however, the authors provide tools that might help with it).

EPIC (3) models the gene expression of the tumor tissue as the sum of the gene expression profiles from the pure cell types composing the sample and uses a constrained least square optimization to perform the deconvolution.

ESTIMATE (4) first pre-determines genes that are likely to define the unique expression patterns of the stromal cells (stromal signature) and the infiltrating immune cells (immune signature). It then uses single-sample Gene Set Enrichment Analysis (ssGSEA) (5) to calculate enrichment scores that serve as a proxy for the presence of infiltrating stromal and immune cells in the tumor samples. Finally, the calculated scores and purity values from a DNA-based method ABSOLUTE (6) are used to train a nonlinear least-squares method to derive the final formula for calculating the tumor purity. The potential downside of this approach is that the genes that are derived for the immune and stromal signature might not capture all of the necessary information required to predict malignant cell proportions. Additionally, ESTIMATE is trained on the estimates of ABSOLUTE, which was one of the earlier methods and might not provide the most accurate genomic-based estimates.

DeMixT (7) models the observed gene expression signal $Y_{ig}$ as a sum of its constituent components: $Y_{ig} = \pi_{1,i}N_{1,ig} + \pi_{2,i}N_{2,ig} + \left(1 - \pi_{1,i} - \pi_{2,i}\right)T_{ig}$ for each gene $g$ and each sample $i$. $\pi_{1,i}$ and $\pi_{2,i}$ are cell proportions, $N_1$, $N_2$ and $T$ are gene expression values within respective cell types. $N_1$- and $N_2$-components are estimated from the available reference samples (normal samples provided as an input), while $T_{ig}$ is derived via inference over a probabilistic model. However, the method's prior probabilistic assumptions are still approximations of the underlying biology, which might introduce additional bias.

LinSeed (8) constructs a collinearity network of genes to determine significantly mutually linear features and keep only them for a later simplex-based deconvolution approach. However, if cell types are closely related to each other, the simplex approach might have trouble distinguishing them as distinct corners (they would be simply too close in the sample vector space). Moreover, our tests have shown that LinSeed's deconvolved cell type proportions are not constrained to sum-to-one, and can sometimes exceed reasonable values (be $> 1$ or $< 0$).

DeconRNASeq (9) uses quadratic programming to solve the weighted non-negative least squares problem $X = AS$ for the cell type proportion matrix $A$, where $X$ is the input gene expression matrix and $S$ is a cell type-specific expression signature matrix. Similar to the CIBERSORTx, the method's performance highly depends on the quality of the signature matrix provided.

In short, while the existing methods employ novel techniques and often use valid assumptions about the deconvolution process, there still are several potential areas for improvement, namely their low correlations with DNA-based estimates, potential bias in the feature selection approach, or being overly- or under-specialized for different cancer types.

**Running transcriptomics-based approaches**

Unless explicitly stated otherwise, the packages below were run in R environment version >= 3.4 (Supplementary code, R). For all the gene expression matrices below, only protein-coding genes were left for the downstream analysis. The gene ids were used in HGNC nomenclature.

CIBERSORTx was run using the web interface available at https://cibersortx.stanford.edu/. The analysis module was selected to be "Impute Cell Fractions", "Custom" mode and "RNA-seq" input data. The NSCLC signature matrix used for imputation was taken from CIBERSORTx's paper (2) supplementary 2l. Mixture files were used in linear space (in TPM values when available, otherwise FPKM) and formatted according to the instructions provided on the website. Batch correction was run in B-mode with no GEP. Quantile normalization was disabled. 100 permutations were used for statistical analysis. "EPCAM" column was taken as tumor purity.

DeMixT (https://github.com/wwylab/DeMixT) was run on gene expression matrices in linear counts space. As the DeMixT package required tumor and normal counts (not necessarily matched), it was run only on datasets that had both available. Additionally, the counts matrices were quartile-normalized and the genes where the total sum of values across all samples was <1 were discarded. As DeMixT seemed to predict the stromal component, the purity was computed as (1-DeMixT predictions).

EPIC (https://github.com/GfellerLab/EPIC) was run on gene expression matrices in linear normalized values space (TPM or FPKM). Purity was taken as the *'otherCells'* component of the resulting *cellFractions.*

ESTIMATE (https://rdrr.io/rforge/estimate/) was run on gene expression matrices in linear normalized values space (TPM or FPKM). For the estimateScore function the *platform* parameter was chosen to be "*affymetrix*".

LinSeed (https://github.com/ctlab/LinSeed) was run on gene expression matrices in linear values space (counts if they were available, or TPM or FPKM if not). Additionally, the matrices were normalized sample-wise so that the samples would have the same sum. RPL/RPS genes were removed from the matrix. *LinseedObject* function was run with *topGenes=10000*, the rest of the functions were run in a 2-component mode according to the instructions provided by the authors in their GitHub repository https://github.com/ctlab/LinSeed. As LinSeed does not explicitly state which of the deconvolved components represents the cancer cells' proportion, the component that had the best Pearson's correlation with DNA-based purities was taken. Additionally, since it appeared that sometimes LinSeed might be predicting the stromal cell proportion instead of purity, if the 1-(LinSeed predictions) had better correlations, that was taken as predicted purity instead.

DeconRNASeq (https://doi.org/doi:10.18129/B9.bioc.DeconRNASeq) was run on gene expression matrices in linear normalized values space (TPM or FPKM), CIBERSORTx's NSCLC matrix was used as a signature. The "EPCAM" column was taken as tumor purity.

There were other methods that we found highly relevant to our work but were not able to include in the benchmark.

Gbm.ensemble is a similar supevised method by Li et al. (10) built as an ensemble of XGBoost models trained on ABSOLUTE tumor purities. Unfortunately, we were not able to run the method following the instructions provided by the authors (https://github.com/yuanyuanli66/gbm.ensemble).

Koo and Rhee (11) built several machine learning models using consensus purity estimates from the literature (CPE values from the "Systematic pan-cancer analysis of tumour purity" by Aran et al. (12)). However, the Github page of their work (https://github.com/BonilKoo/ML_purity) does not present a working method and so this approach could not be added to the benchmark.

DeClust by Wang et al. (13) is a method for reference profile-free deconvolution method to infer cancer cell-intrinsic subtypes, that could potentially be used to estimate cancer cell proportion in the tissues. However, we were not able to run the method for the tumor purity estimation purposes using the code provided by the authors (https://github.com/integrativenetworkbiology/DeClust).

##### Selected gene set enrichment analysis

The gene set analysis was run in Python using the "GSEApy" package (https://github.com/zqfang/GSEApy). The names of 158 selected genes were converted into HGNC nomenclature and used as an input to the enrichr function. Background genes were set to be 9554 significantly expressed autosomal genes in TCGA. The hallmark gene set used in the enrichment analysis was downloaded from the MSigDB collection (http://www.gsea-msigdb.org/gsea/msigdb/collections.jsp, set H). For each gene, Pearson correlation was computed between its expression and DNA-based tumor purity in each of the 20 cancer types in the TCGA train set, and a mean of it was taken. Top 10 enriched pathways by Benjamini-Hochberg adjusted p-value were computed for 4 gene sets: full feature set of 158 genes, top 30% genes by their mean expression-purity correlation per cancer type, bottom 30% genes by their mean expression-purity correlation per cancer type, and, finally, the genes in the 30-70% range by their mean expression-purity correlation per cancer type.

| Abbreviation | Full name | Number of samples (in train set / in test set) | Median tumor purity (in train set / in test set) |
|---|---|---|---|
| BRCA | Breast invasive carcinoma | 1060 (848 / 212) | 0.56 (0.56 / 0.55) |
| LUAD | Lung adenocarcinoma | 508 (406 / 102) | 0.42 (0.42 / 0.44) |
| LGG | Brain lower grade glioma | 506 (405 / 101) | 0.66 (0.67 / 0.62) |
| HNSC | Head and neck squamous cell carcinoma | 494 (395 / 99) | 0.47 (0.47 / 0.47) |
| PRAD | Prostate adenocarcinoma | 489 (391 / 98) | 0.48 (0.48 / 0.5) |
| THCA | Thyroid carcinoma | 485 (388 / 97) | 0.52 (0.52 / 0.54) |
| LUSC | Lung squamous cell carcinoma | 482 (385 / 97) | 0.49 (0.49 / 0.48) |
| STAD | Stomach adenocarcinoma | 410 (328 / 82) | 0.47 (0.46 / 0.47) |
| BLCA | Bladder urothelial carcinoma | 403 (322 / 81) | 0.56 (0.57 / 0.56) |
| KIRC | Kidney renal clear cell carcinoma | 377 (302 / 75) | 0.51 (0.51 / 0.53) |
| CRC | Combined: colon adenocarcinoma (COAD) and rectum adenocarcinoma (READ) | 367 (294 / 73) | 0.62 (0.62 / 0.62) |
| SKCM | Skin cutaneous melanoma | 365 (292 / 73) | 0.64 (0.66 / 0.57) |
| LIHC | Liver hepatocellular carcinoma | 360 (288 / 72) | 0.66 (0.66 / 0.67) |
| OV | Ovarian serous cystadenocarcinoma | 301 (241 / 60) | 0.72 (0.72 / 0.76) |
| CESC | Cervical squamous cell carcinoma and endocervical adenocarcinoma | 292 (234 / 58) | 0.61 (0.6 / 0.63) |
| KIRP | Kidney renal papillary cell carcinoma | 284 (227 / 57) | 0.66 (0.66 / 0.61) |
| ESCA | Esophageal carcinoma | 180 (144 / 36) | 0.56 (0.54 / 0.59) |
| UCEC | Uterine corpus endometrial carcinoma | 179 (143 / 36) | 0.67 (0.66 / 0.67) |
| PAAD | Pancreatic adenocarcinoma | 176 (141 / 35) | 0.35 (0.36 / 0.35) |
| GBM | Glioblastoma multiforme | 146 (117 / 29) | 0.72 (0.73 / 0.72) |

**Suppl. Table S1: TCGA dataset description and train/test set distribution.** TCGA dataset used in this study was compiled from datasets of 20 solid cancer types, resulting in 7864 samples (6291 in the train set, 1573 in the test set). Median purity is rounded to 2 digits after the decimal point.

| Cohort name | Cancer type | Num. of samples | Median purity | Purity estimation method |
|---|---|---|---|---|
| TCGA (test set) | 20 solid cancer types | 1573 | 0.56 | Consensus (ABSOLUTE, ASCAT, PurBayes, AbsCNSeq) |
| Chen et al. | Lung | 172 | 0.45 | Sequenza |
| Chua et al. | Lung | 64 | 0.6 | Consensus (theta2, TitanCNA, PurBayes, AbsCNSeq) |
| Joanito et al. | Colorectal | 153 | 0.62 | Consensus (theta2, TitanCNA, PurBayes, AbsCNSeq) |
| TCGA-CRC+ | Colorectal | 243 | 0.7 | ABSOLUTE |
| TCGA-UCEC+ | Uterine | 353 | 0.76 | ABSOLUTE |
| TCGA-PCPG | Paraganglioma | 164 | 0.74 | ABSOLUTE |
| TCGA-TGCT | Testicular | 155 | 0.6 | ABSOLUTE |

**Suppl. Table S2:** Independent datasets descriptions and purity statistics. Median purity is rounded to 2 digits after the decimal point.

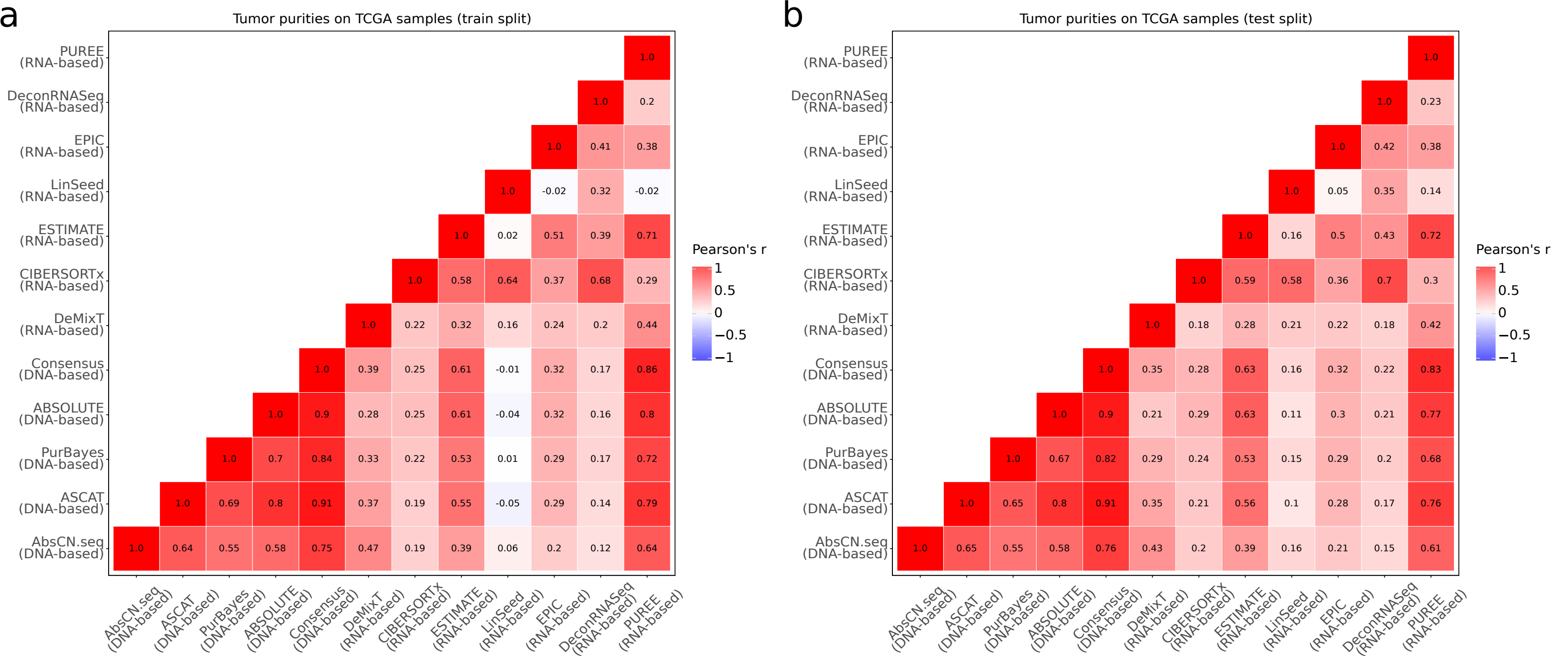


**Suppl. Figure S1: Agreement of the genomics- and transcriptomics-based purity estimation methods.** Pairwise correlations of tumor purity estimates were computed separately for **a)** TCGA train (6291 samples) and **b)** TCGA test (1573 samples) splits of the dataset to account for the machine learning training procedure of PUREE.

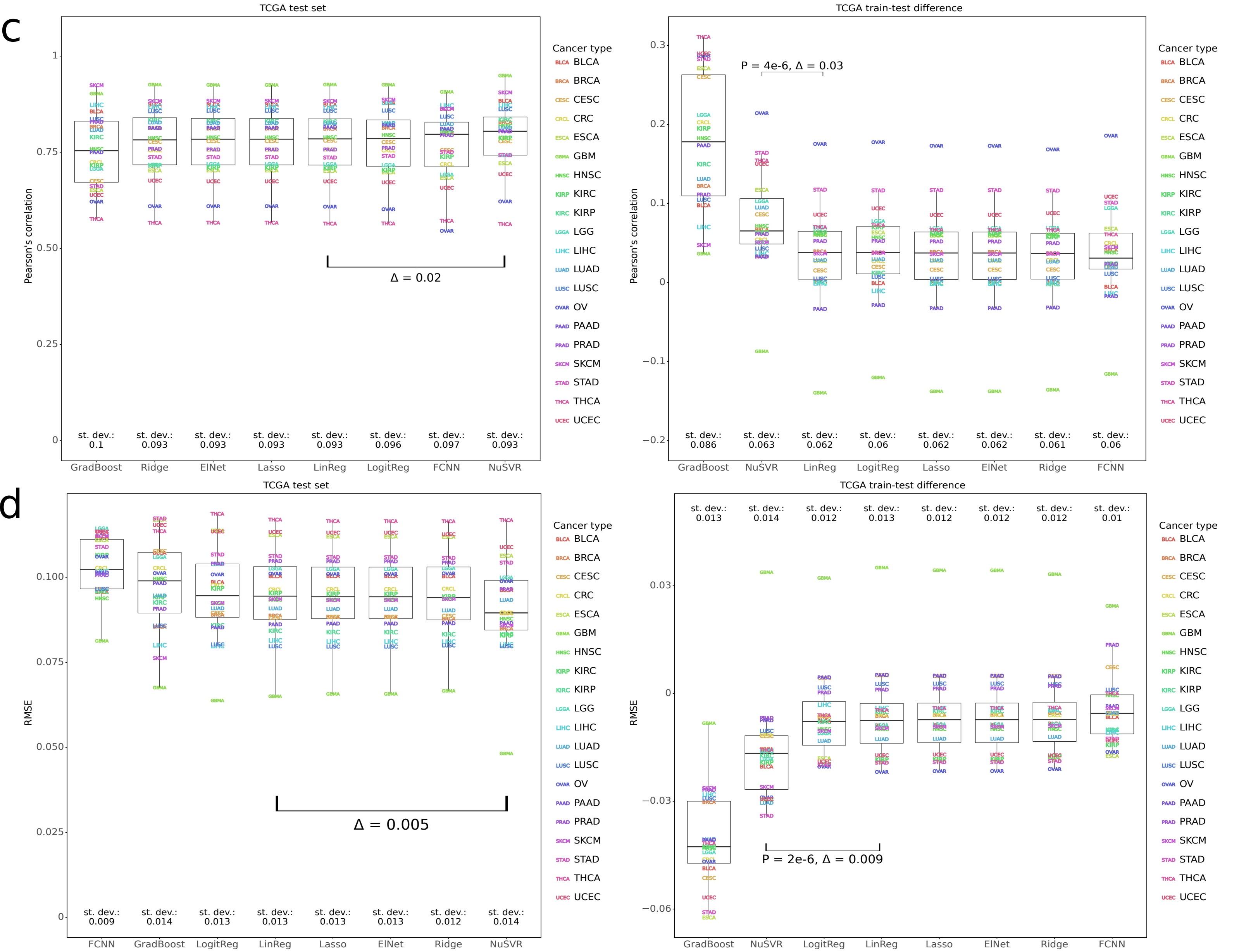


**Suppl. Figure S2: Exploring the range of machine learning methods for the PUREE approach. a,b)** models trained on the full set of 9554 features (a - Pearson's correlation, b - RMSE); lasso was chosen as the best for the full feature set due to good performance and low train-test difference in performance. **c,d)** models trained on the reduced feature set of 158 features (c - Pearson's correlation, d - RMSE); linear regression was chosen over NuSVR due to comparable performance, lower train-test difference in performance and simplicity. Each column depicts performance on the TCGA test set and the difference in performance between train and test sets respectively. Gradient Boosting, NuSVR, Ridge, Lasso and Elastic Net models were trained using 5-fold cross-validation hyperparameter selection procedure. Numbers next to boxplots indicate standard deviation. Delta between boxplots medians and Wilcoxon signed-rank test (two-tailed) shown for two best models in each plot (NuSVR and Lasso for 10K and NuSVR and Linear Regression for the reduced set).

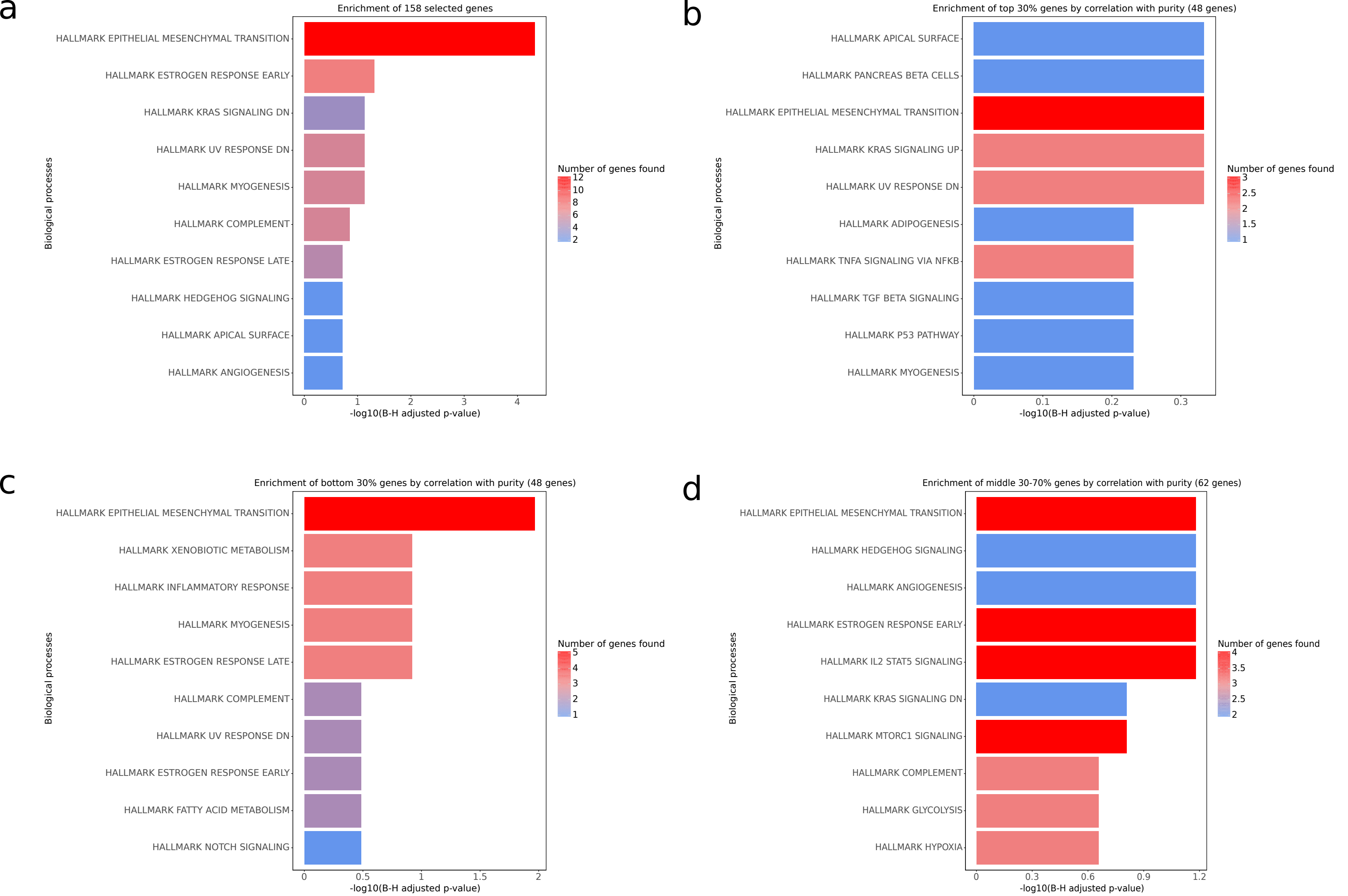


**Suppl. Figure S3: Hallmark pathway analysis of** 158 **genes used in the final model.** Only the top 10 pathways ranked by Benjamini-Hochberg adjusted P-value shown. **a)** enrichment analysis of 158 selected genes; **b)** enrichment analysis of top 30% genes by expression-purity correlations (correlation above 0.05); **c)** enrichment analysis of bottom 30% genes by expression-purity correlations (correlation below -0.14); **d)** enrichment of genes whose expression-purity correlations lie in the 30-70% ranking interval (correlation between 0.05 and -0.14). Enrichment analysis performed with GSEApy package in Python. Expression-purity correlations calculated using averages of mean correlations per cancer type. Pearson's correlation used.

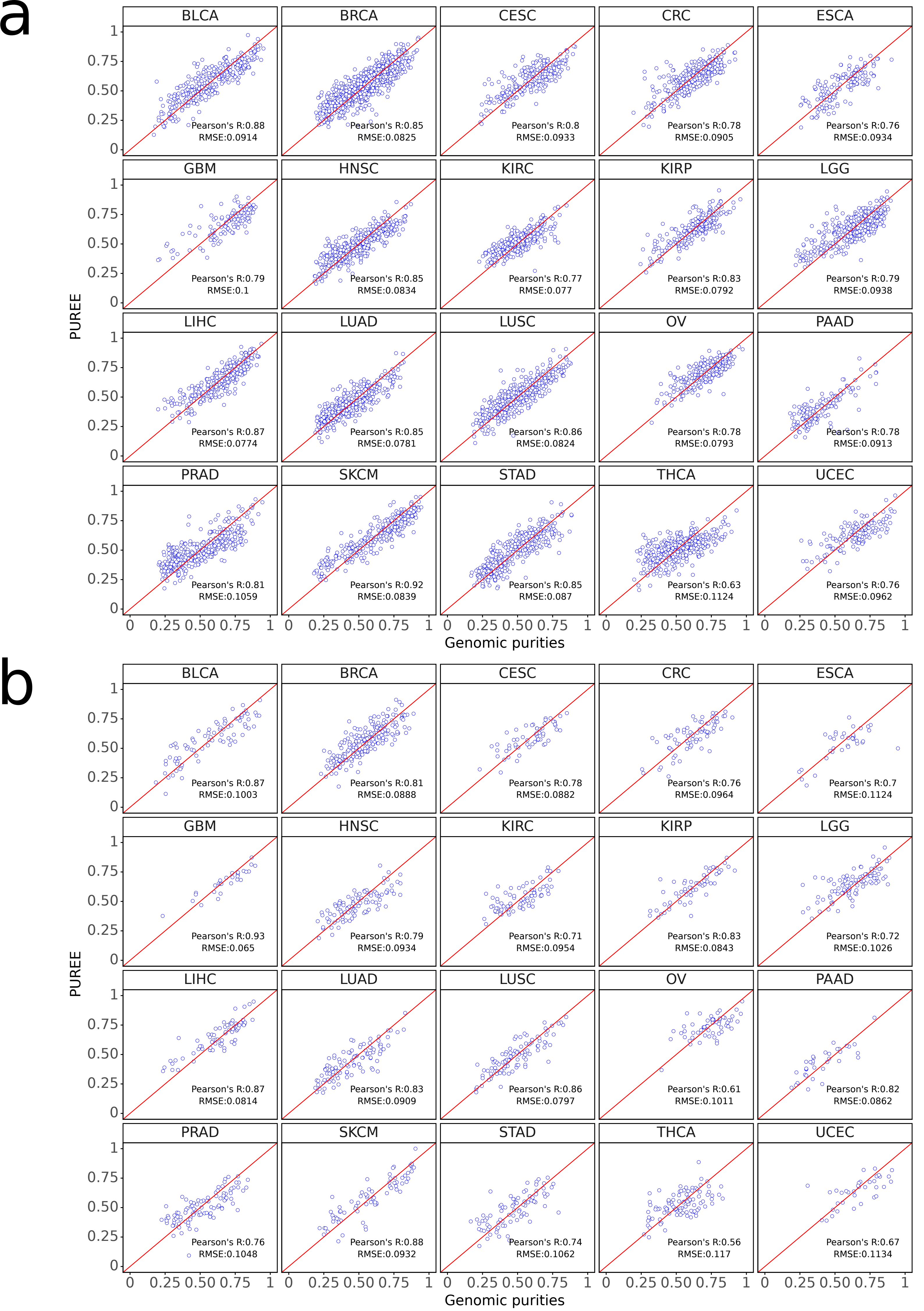


**Suppl. Figure S4: Raw data of purity predictions of PUREE across 20 cancer types from TCGA. a)** TCGA train split (6291 samples). **b)** TCGA test split (1573 samples).

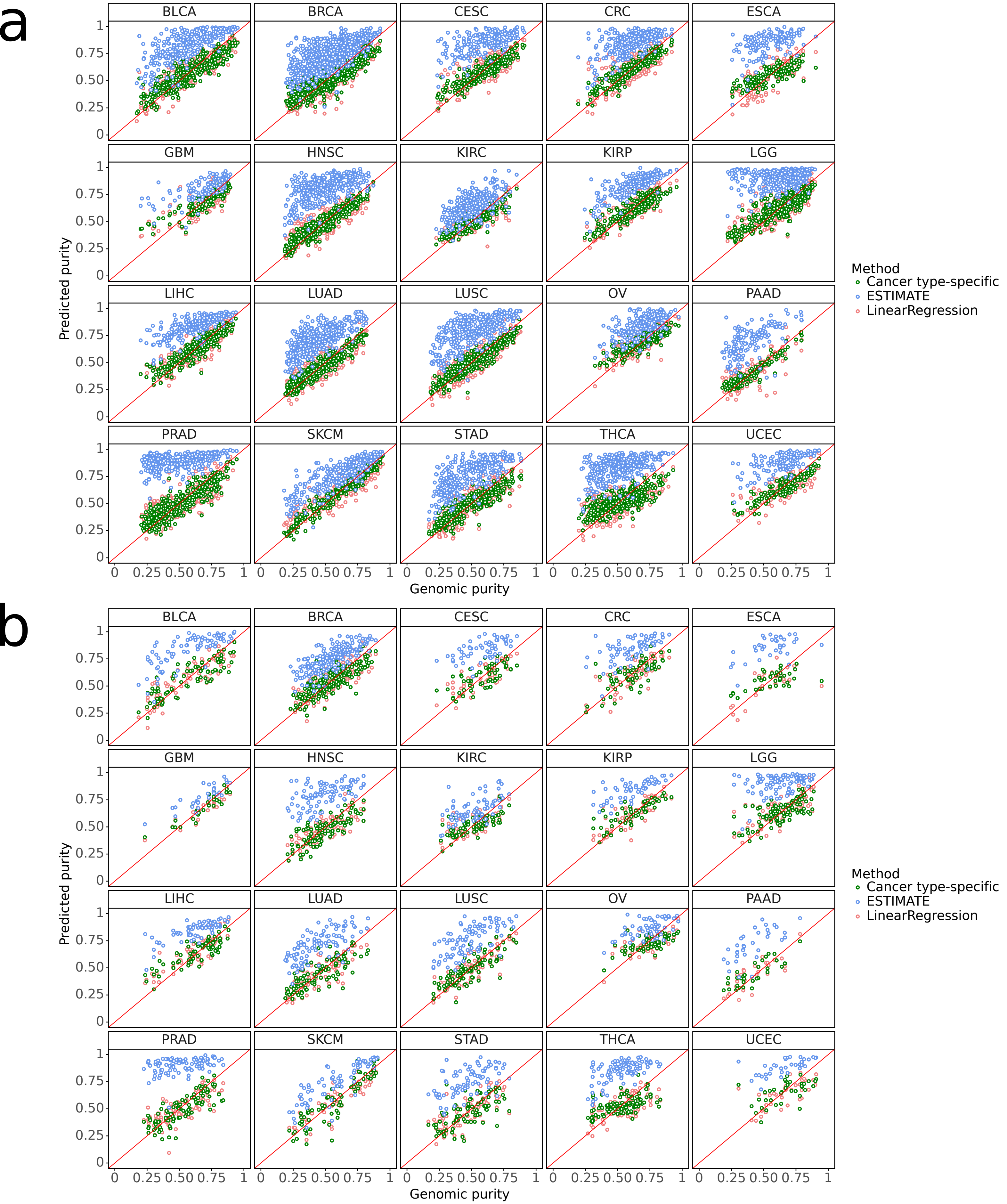


**Suppl. Figure S5: Raw data of purity predictions of cancer-specific models, PUREE and ESTIMATE across 20 cancer types from TCGA** on **a)** TCGA train set (6291 samples) and **b)** TCGA test set (1573 samples).

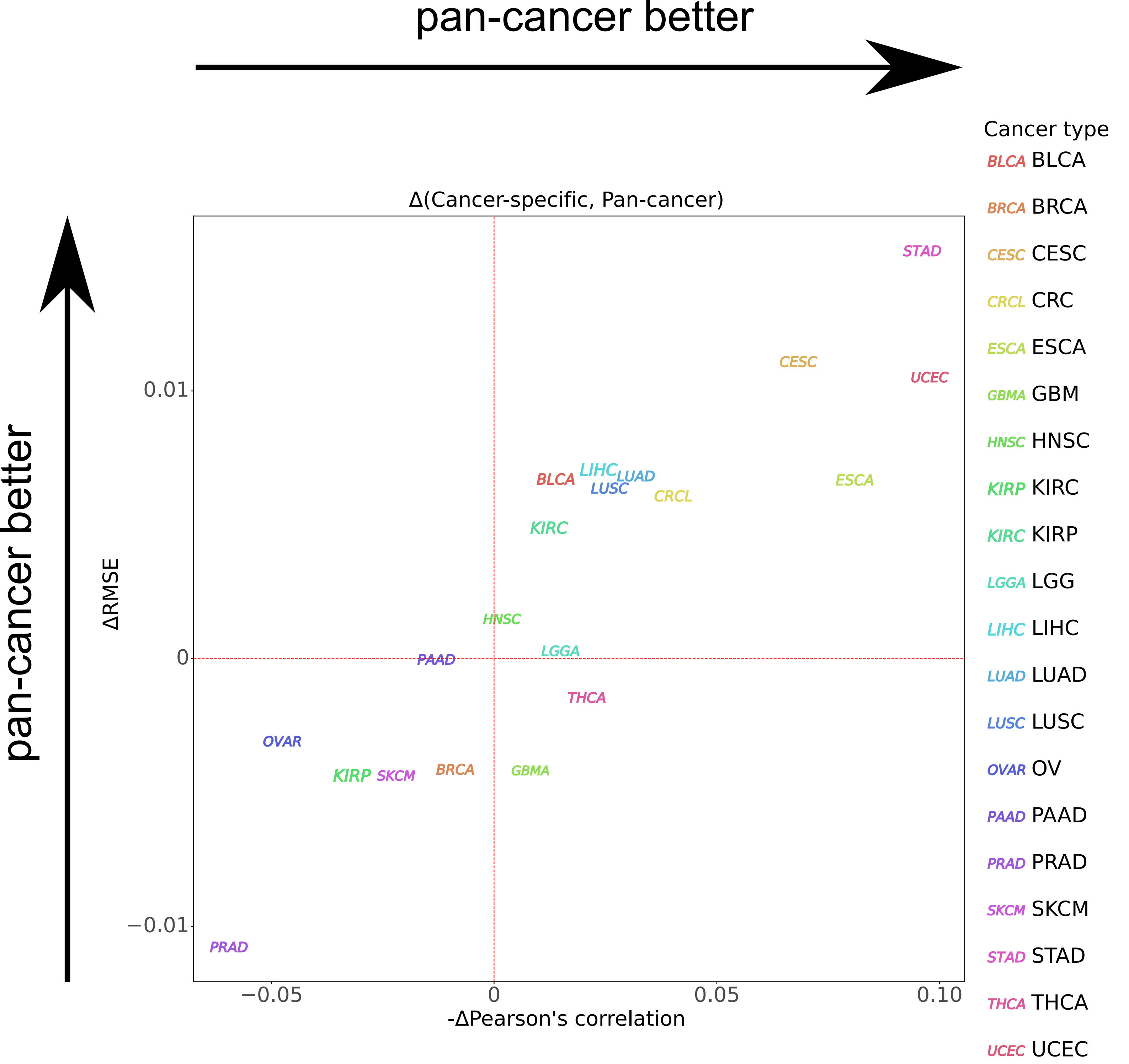


**Suppl. Figure S6: Cancer type-specific vs pan-cancer test.** Delta between average metrics per cancer type between the pancancer model (PUREE) and the cancer type-specific ones.

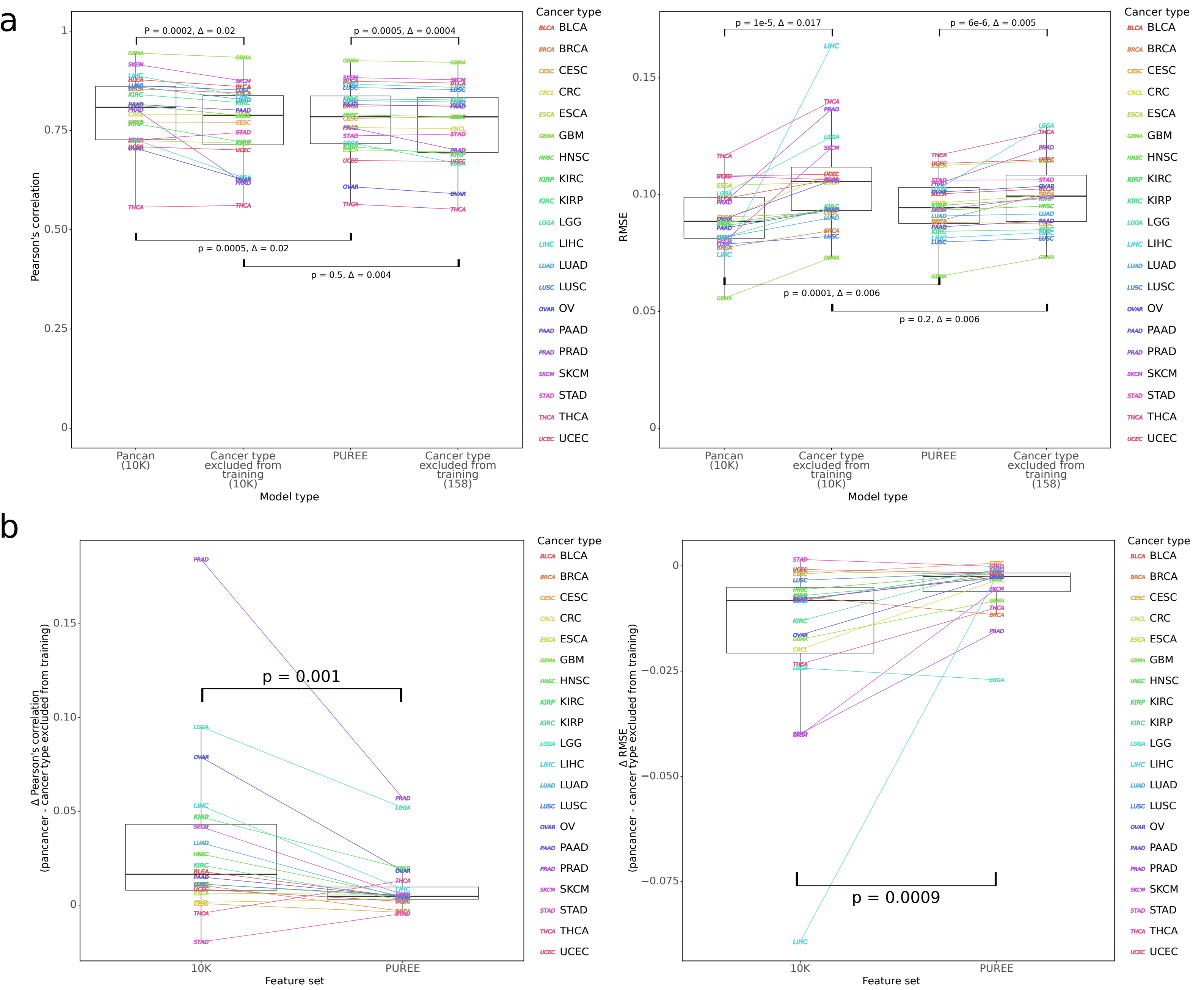


**Suppl. Figure S7: One cancer type excluded from training test. a)** Means of Pearson's correlation (left) and RMSE (right) for 20 cancer types shown for one cancer type excluded-test for the model built on 10K features versus PUREE. **b)** Differences of means of Pearson's correlation (left) and RMSE (right) for 20 cancer types per feature set (10K versus PUREE), demonstrating the effect of using the reduced feature set. P-values calculated using the Wilcoxon signed-rank test (two-tailed). Delta provided is the difference between medians of metrics across all cancer types.

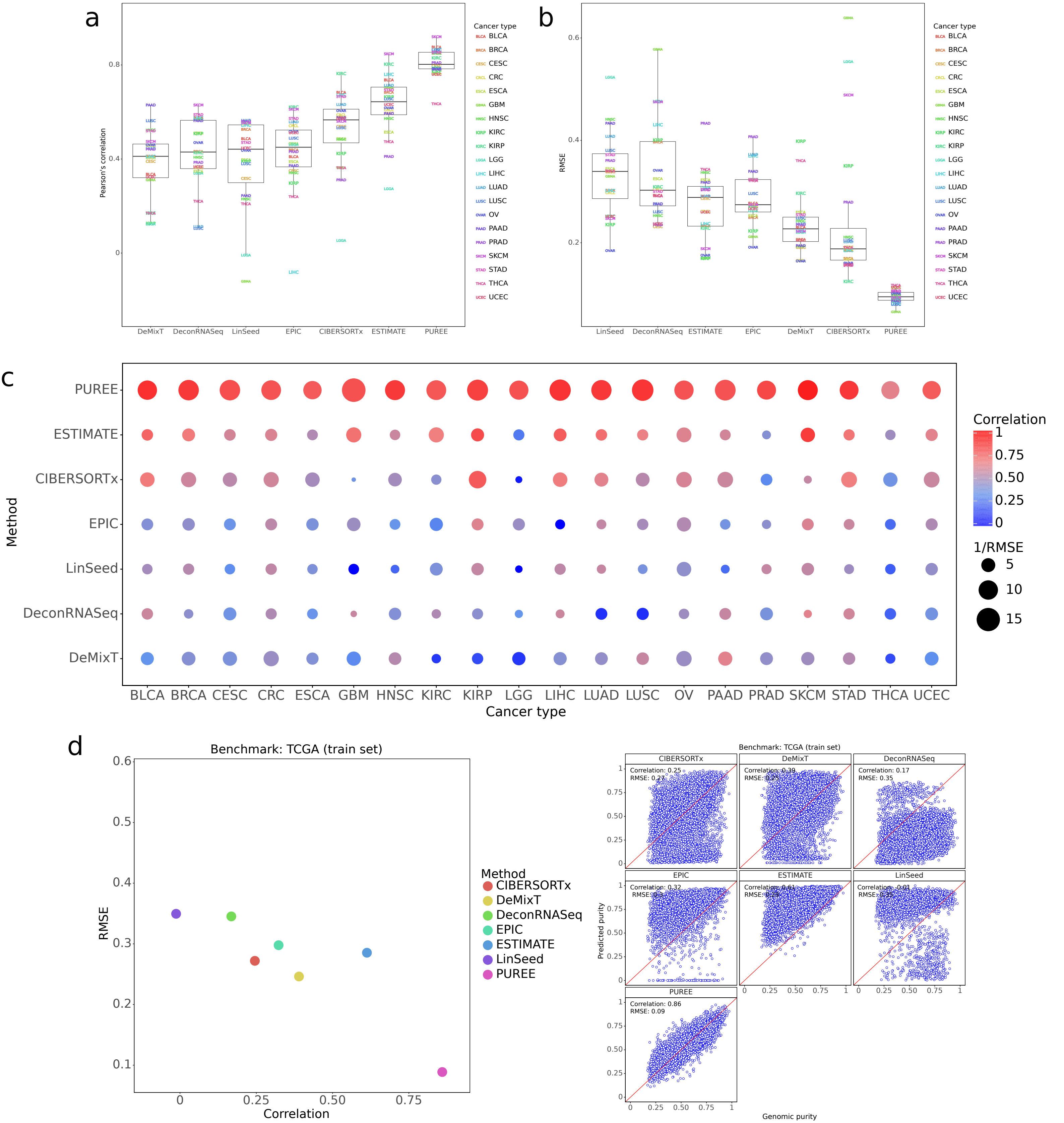


**Suppl. Figure S8: RMSE and correlation on TCGA train set. a,b,c)** PUREE's and 6 other transcriptomics-based deconvolution methods' performance on TCGA train set (6291 samples), shown across 20 cancer types; mean Pearson's correlation (a) and RMSE (b) separately, and aggregated (c). **d)** Raw data of purity predictions for TCGA train set (left: aggregated correlation-RMSE plot, right: raw data).

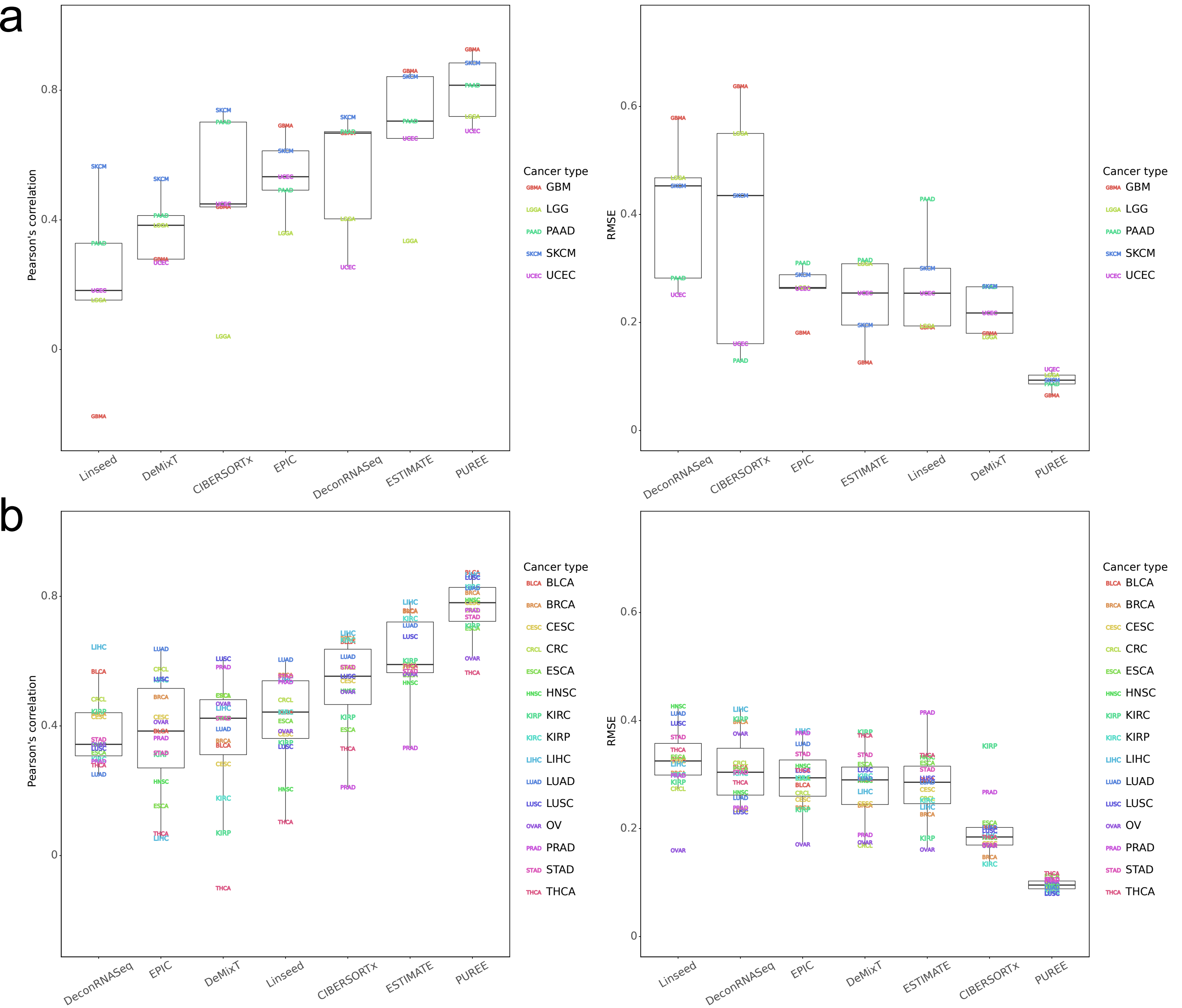


**Supplementary Figure S9: Performance of PUREE and 6 other transcriptomics-based methods on TCGA test set for selected cancer types.** Mean Pearson's correlation and RMSE of methods per cancer type when compared with genomic tumor purity estimates on the TCGA test data split (1573 samples); a) data shown for cancer types with potentially high levels of dissimilar stroma (GBM, LGG, PRAD, SKCM, UCEC), b) data shown for other 15 cancer types.

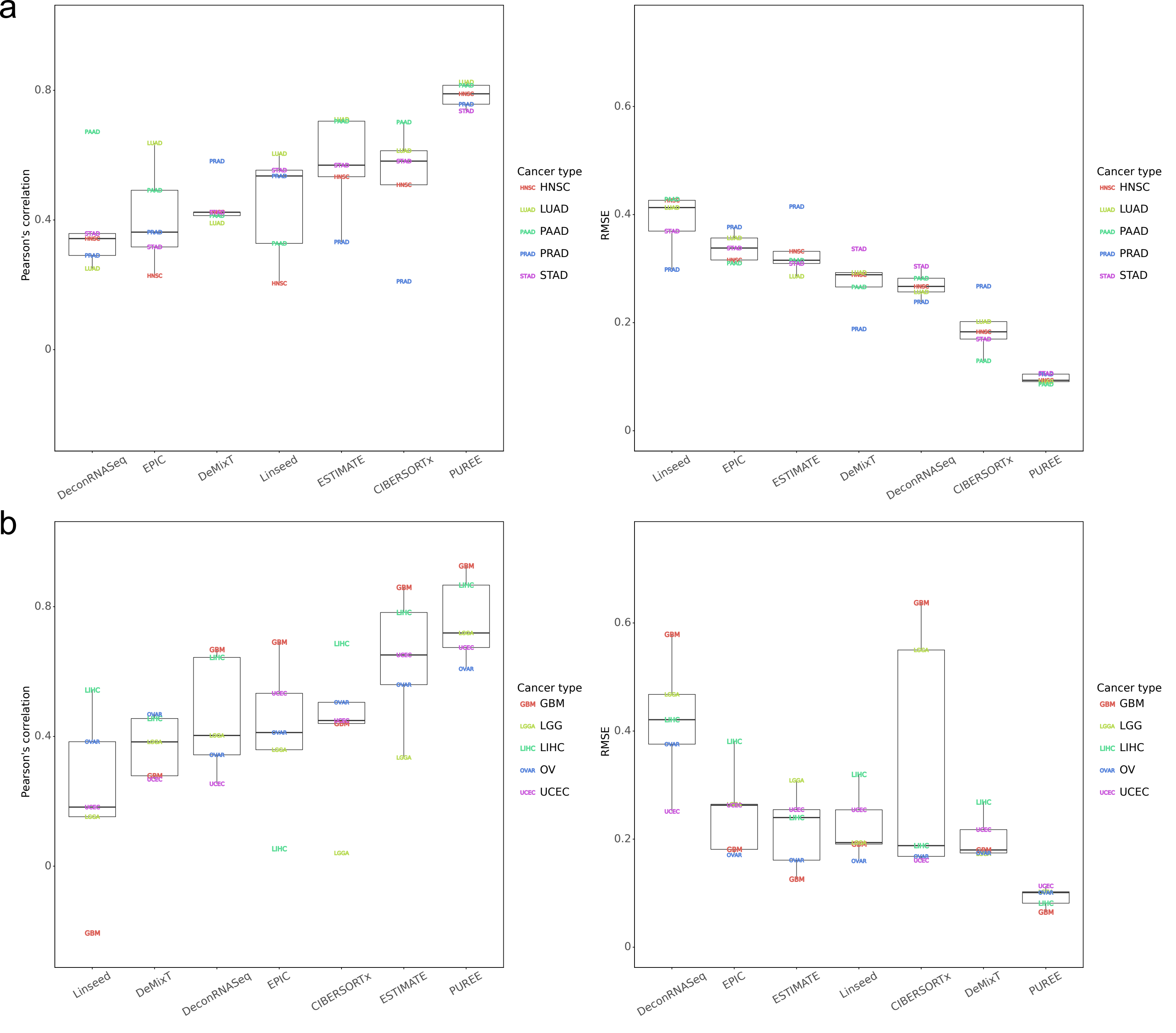


**Supplementary Figure S10: Performance of PUREE and 6 other transcriptomics-based methods on TCGA test set for selected cancer types with different median purities.** Mean Pearson's correlation and RMSE of methods per cancer type when compared with genomic tumor purity estimates on the TCGA test data split (1573 samples); a) cancer types with the lowest median consensus tumor purity (HNSC, LUAD, PAAD, PRAD, STAD), b) cancer types with the highest median consensus tumor purity (GBM, LGG, LIHC, OV, UCEC).

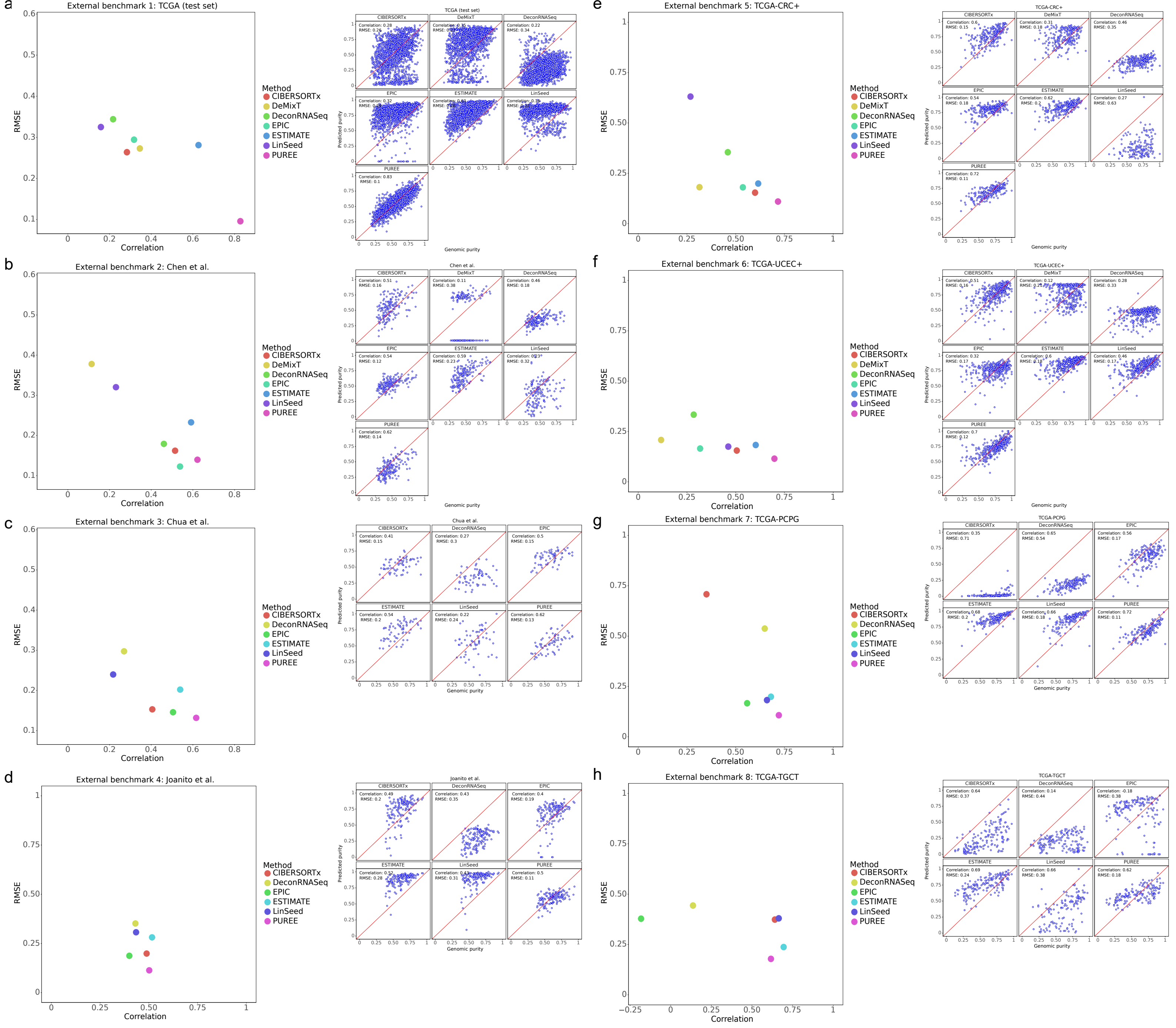


**Suppl. Figure S11: External benchmarks' for PUREE's and 6 other transcriptomics-based methods for tumor purity prediction.** Aggregated correlation-RMSE performance plots (left) and raw data (right) for a) the test portion of TCGA (1573 samples); b) Chen et al. cohort, lung cancer (172 samples); c) Chua et al. cohort, lung cancer (64 samples); d) Joanito et al. cohort, colorectal cancer (153 samples); e) TCGA-CRC+ (243 samples), colorectal cancers; f) TCGA-UCEC+ (353 samples), uterine endometrial cancer; g) TCGA-PCPG, pheochromocytoma and paraganglioma (164 samples) and h) TCGA-TGCT (155 samples), testicular cancer. Pearson's r correlation coefficient shown.

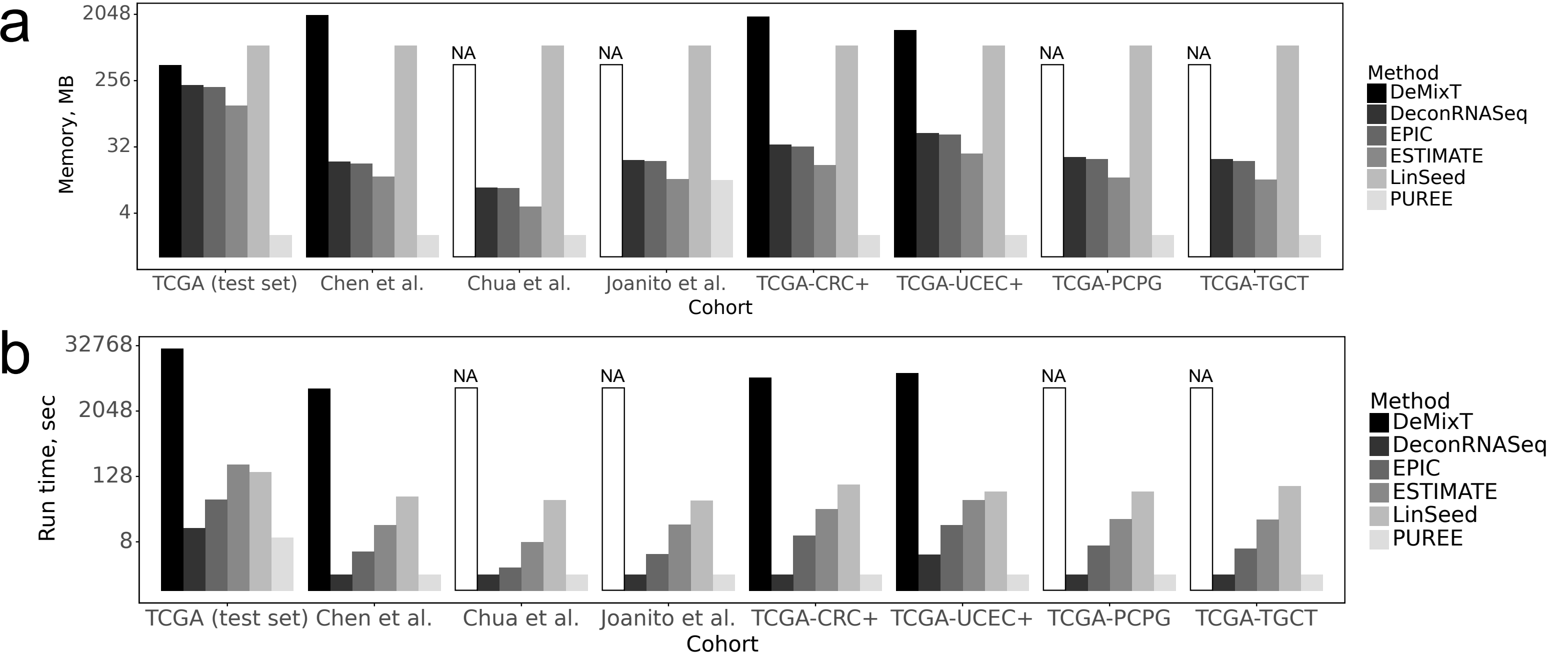


**Suppl. Figure S12: Time-memory benchmark on TCGA test split and 7 external cohorts. a)** Peak memory used by the prediction function. **b)** Execution time of the prediction function. The time and memory to load the data and save the results were not taken into account. DeMixT could not be run on the Chua et al., Joanito et al., PCPG and TGCT cohorts due to the absence of normal samples there.

a

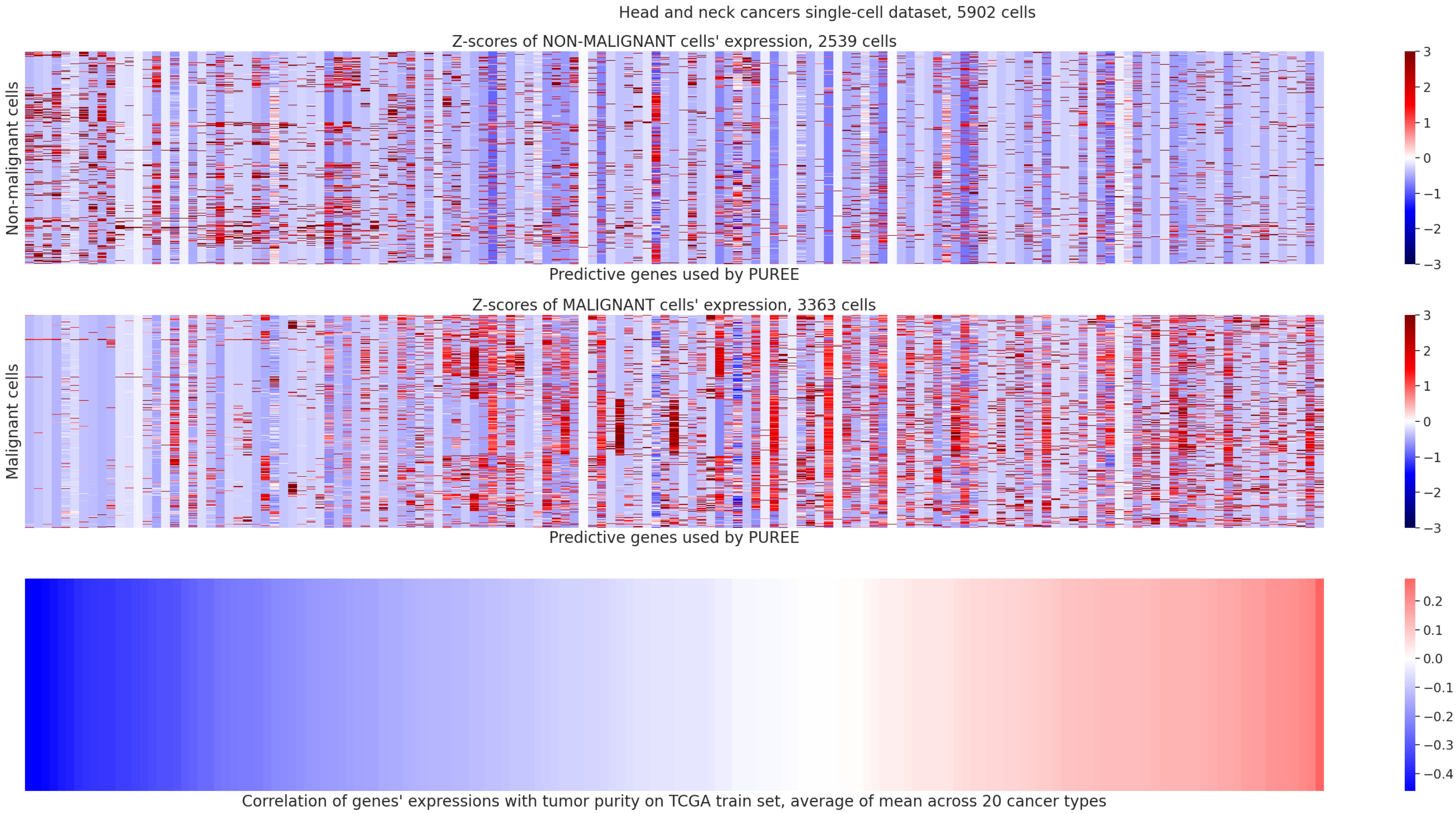


b

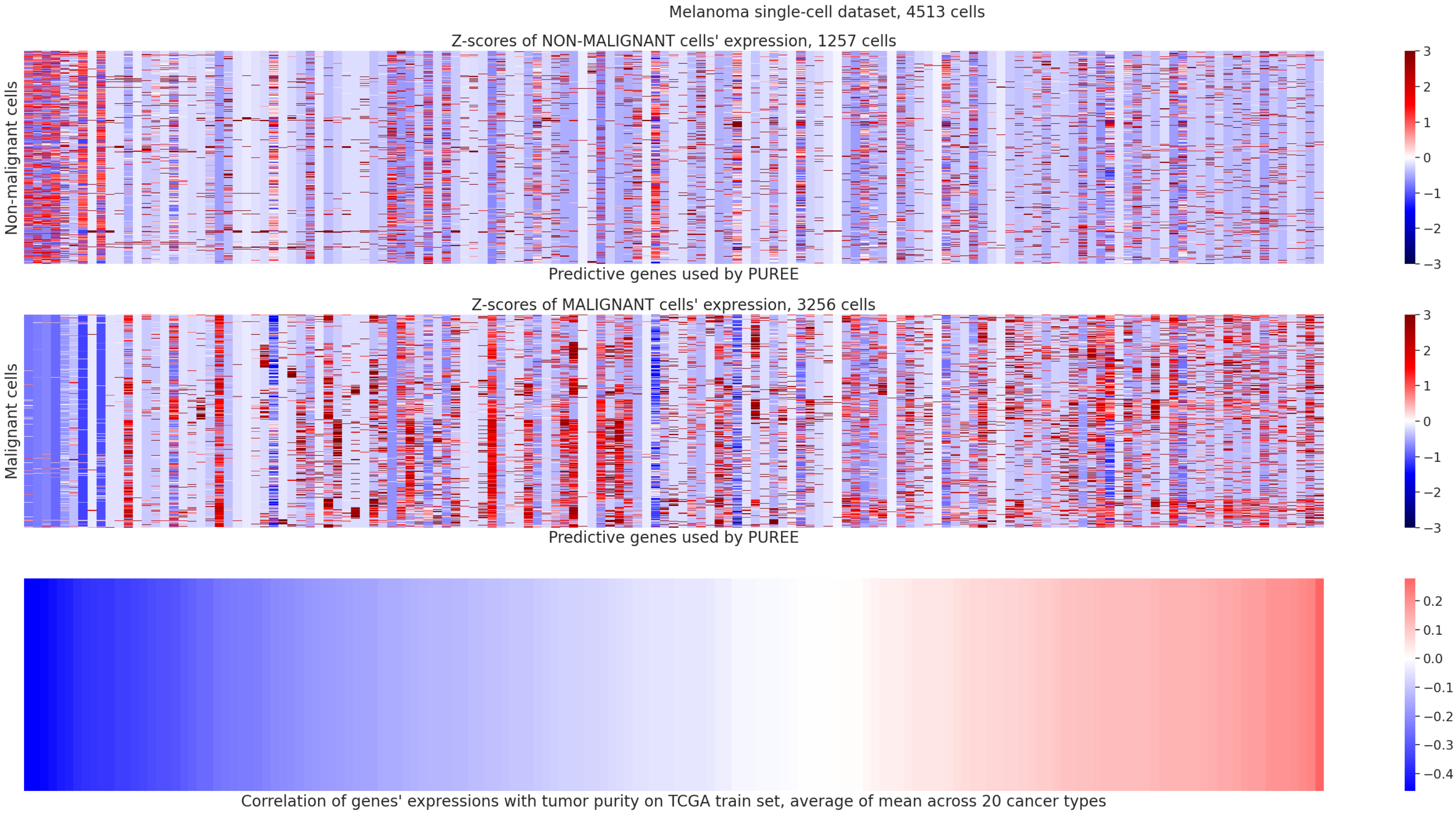


**Suppl. Figure S13: Heatmaps of PUREE's genes' expressions in single-cell RNA-seq data. a)** Puram et al., 2017, head and neck cancers; **b)** Tirosh et al., 2016, melanoma. Genes (columns) sorted by mean correlations of their expressions with genomic-based tumor purity on TCGA train set. Correlations were computed as means for each cancer type first and then averaged across all cancer types. Genes present in the 158 features of PUREE but missing from the single cell data were dropped, which resulted in 143 genes left in both datasets.
